# A reverse-transcription loop-mediated isothermal amplification (RT-LAMP) assay for the rapid colorimetric detection of pepper mild mottle virus (PMMoV)

**DOI:** 10.1101/2025.05.14.654098

**Authors:** Anthony J Gross, Salvador Lopez, Alexandra Rogers, Scott Adkins, Mya Breitbart

**Affiliations:** College of Marine Science, University of South Florida, Saint Petersburg, FL USA; USDA-ARS, US Horticultural Research Laboratory, Fort Pierce, FL, USA

**Author notes:** Corresponding author: Mya Breitbart.

**Keywords:** Pepper mild mottle virus (PMMoV), Reverse-transcription loop-mediated isothermal amplification (RT-LAMP), *Tobamovirus capsici*, Virus detection

## Abstract

Pepper mild mottle virus (*Tobamovirus capsica*, PMMoV) is a plant virus in the genus *Tobamovirus* that infects peppers and other members of the family *Solanaceae*. The virus is transmitted mechanically, poses a significant threat to crops globally, and is one of the most abundant viruses found in human feces and wastewater. Two colorimetric reverse-transcription loop-mediated isothermal amplification (RT-LAMP) assays were developed to detect PMMoV, one targeting the RNA-dependent RNA-polymerase (PMMoV_RdRp) and the other targeting the coat protein (PMMoV_CP). Synthetic gBlock positive controls were used to determine the detection limit of each assay. PMMoV_RdRp detected PMMoV at concentrations greater than or equal to 100 copies/μL, the same sensitivity as the RT-qPCR assay for this gene. In contrast, the detection limit of the PMMoV_CP RT-LAMP assay was an order of magnitude greater. Both assays were specific to PMMoV and did not amplify plant host tissue or related tobamoviruses. Since these RT-LAMP assays do not require specialized laboratory equipment and yield positive results within 20-30 minutes, they are advantageous for point-of-use testing. Overall, the RT-LAMP assays described here are sensitive, specific, and more rapid than existing methods for PMMoV detection and quantification and thus have potential widespread applications for agriculture, wastewater treatment assessment, recreational water quality testing, and food safety.

**Highlights:**

- RT-LAMP assays were developed for the detection of the PMMoV RdRp and CP genes
- The RdRp assay matches the detection limit of the established PMMoV RT-qPCR assay
- Positive results are obtained within 20-30 minutes from the reaction start
- These RT-LAMP assays are specific to PMMoV and do not amplify related viruses
- These assays are applicable for PMMoV detection across diverse scientific fields

## 1. Introduction

Pepper mild mottle virus (*Tobamovirus capsica*, PMMoV) is a plant virus in the genus *Tobamovirus*, family *Virgaviridae* (Simmonds et al., 2024). PMMoV was first isolated and described by researchers in 1984 from *Capsicum annum* (bell pepper) in Sicily, Italy (Wetter, 1984). PMMoV mainly infects Capsicum spp. but can also infect plants belonging to the families *Chenopodiaceae, Cucurbitaceae, Labiatae, Solanaceae*, and *Plantaginaceae* (Kumari et al., 2023; Wetter, 1984; Zhou et al., 2021). The PMMoV virion is rod-shaped, ~312 nm long, and encapsulates a single copy of the 6.4 kb, positive-sense, single-stranded RNA genome (Kitajima et al., 2018; Wetter, 1984). PMMoV has an isoelectric point of pH 3.2 - 3.8, similar to other tobamoviruses (Shirasaki et al., 2017; Wetter, 1984).

PMMoV is extremely stable and infective in soil, humus, compost, leftover plant debris, and agricultural tools (Ochar et al., 2023). The virus is transmitted via mechanical contact, primarily through infected seeds, making the planting process an opportunity for infection in agricultural settings (Ochar et al., 2023). Owing to its high infectivity, PMMoV is an economically significant virus that causes severe crop losses globally. PMMoV infection is characterized by leaf mosaic, mottling, stunting, fruit deformities, and reduced fruit yields (Adkins et al., 2001; Kumari et al., 2023). PMMoV infection results in an average of 15 - 40% loss in fruit yield (Kim et al., 2010); however, in a single case, an entire pepper field was lost due to the disease (Martínez-Ochoa et al., 2003).

In addition to its worldwide agricultural significance, PMMoV is also one of the most abundant viruses found in human feces and wastewater globally (Eifan et al., 2023; Rosario et al., 2009; Rothman and Whiteson, 2022; Symonds et al., 2018; Zhang et al., 2005). PMMoV dominates the RNA viral community in human feces, present at concentrations of up to 10^9^ copies per gram dry weight of fecal material (Zhang et al., 2005). PMMoV is also abundant in wastewater, where it can be found at concentrations ranging from 1.10 × 10^4^ to 7.00 × 10^6^ copies/mL (Rosario et al., 2009; Rothman and Whiteson, 2022). PMMoV occurrence in human feces is dietary in origin and sourced from eating foods that PMMoV infects and their processed products, such as peppers and hot sauce (Zhang et al., 2005). PMMoV is consistently found at high concentrations in human fecal samples but is generally not detected in other animals, except for sporadic reports of low concentrations in bird and dog feces (Gyawali et al., 2019; Rosario et al., 2009). This enables the use of PMMoV as a viral indicator of human fecal pollution in source tracking of sewage and wastewater contamination in environmental waters (Ahmed et al., 2018; Symonds et al., 2019, 2018, 2016).

Reverse transcription-quantitative PCR (RT-qPCR) is currently the standard detection method for PMMoV in wastewater and plant tissue, although reverse transcription PCR (RT-PCR) followed by sequencing is also used. Using RT-qPCR allows for real-time detection of PMMoV down to concentrations of 100 copies per reaction (Rosario et al., 2009; Symonds et al., 2016). Other detection methods for PMMoV include commercially available serological techniques such as enzyme-linked immunosorbent assay (ELISA) (Adkins et al., 2001) and lateral flow immunoassays.

Loop-mediated isothermal amplification (LAMP), and by extension, reverse-transcriptase loop-mediated isothermal amplification (RT-LAMP), is a method for the rapid, specific, and efficient amplification and detection of nucleic acids at an isothermal temperature (Notomi et al., 2000). Unlike PCR-based assays, RT-LAMP utilizes a set of four to six primers – one forward and one backward outer primer (F3/B3) and one forward inner and one backward inner primer (FIP/BIP). Six-primer LAMP assays also include one forward and one backward loop primer (LF/LB) to accelerate the reaction (Nagamine et al., 2002). Recent research has found that using only one loop primer compared to two reduces the false-positive rate of the reaction by delaying or preventing misamplifications (Alhamid et al., 2023). RT-LAMP assays have been previously developed for other tobamoviruses, such as tobacco mosaic virus (*Tobamovirus tabaci*, TMV) (Liu et al., 2010) and tobacco mild green mosaic virus (*Tobamovirus mititessellati*, TMGMV) (Zhao et al., 2021) and a recent manuscript described a single nucleotide polymorphism-specific duplex P*f*Ago RT-LAMP assay for PMMoV and SARS-CoV-2 (Oh et al., 2024).

## 2. Materials and Methods

### 2.1 Primer Design

For this study, we developed two RT-LAMP assays (one six-primer and one five-primer) for the detection of PMMoV (U.S. Patent Application 19/049,344). RT-LAMP primers were designed using the PrimerExplorerV5 website software (https://primerexplorer.jp/lampv5e/index.html) and ordered from Integrated DNA Technologies (IDT, USA). The PMMoV_RdRp primer set (Table 1) targets the PMMoV RNA-dependent RNA-polymerase (RdRp) gene (positions 1957 - 2156) on the PMMoV genome (Accession # NC_003630.1; Table 2). The PMMoV_CP primer set (Table 1) targets the PMMoV CP gene (positions 5871 - 6091) on the PMMoV genome (Accession # NC_003630.1; Table 2).

**Table 1.**
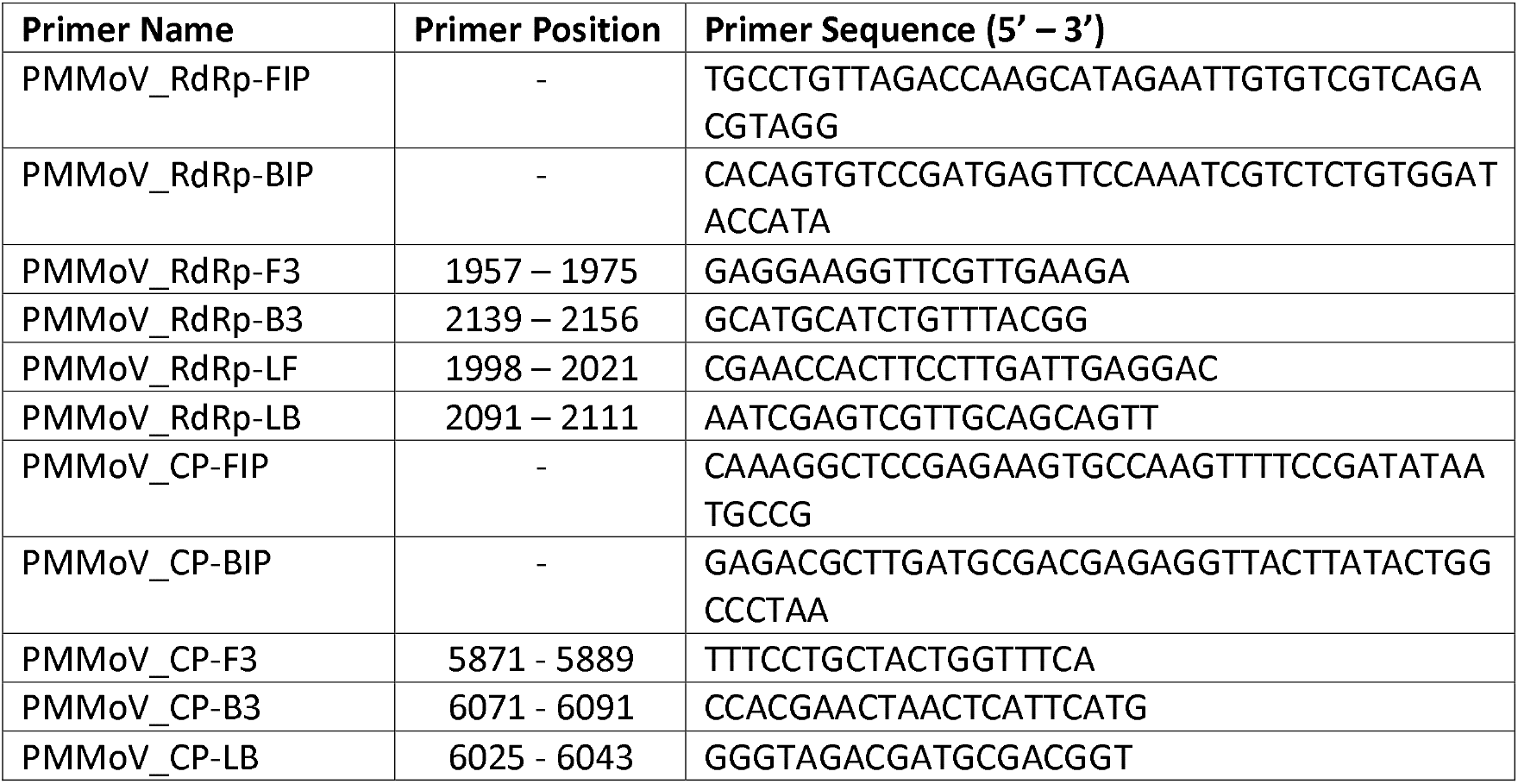
PMMoV_RdRp and PMMoV_CP primers used for the primer mix with respective genome position and sequence.

**Table 2.**
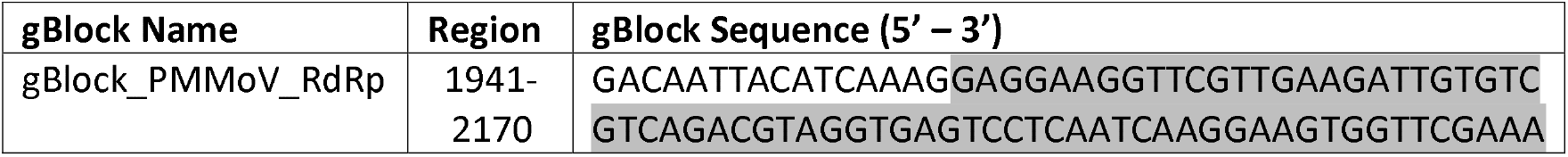
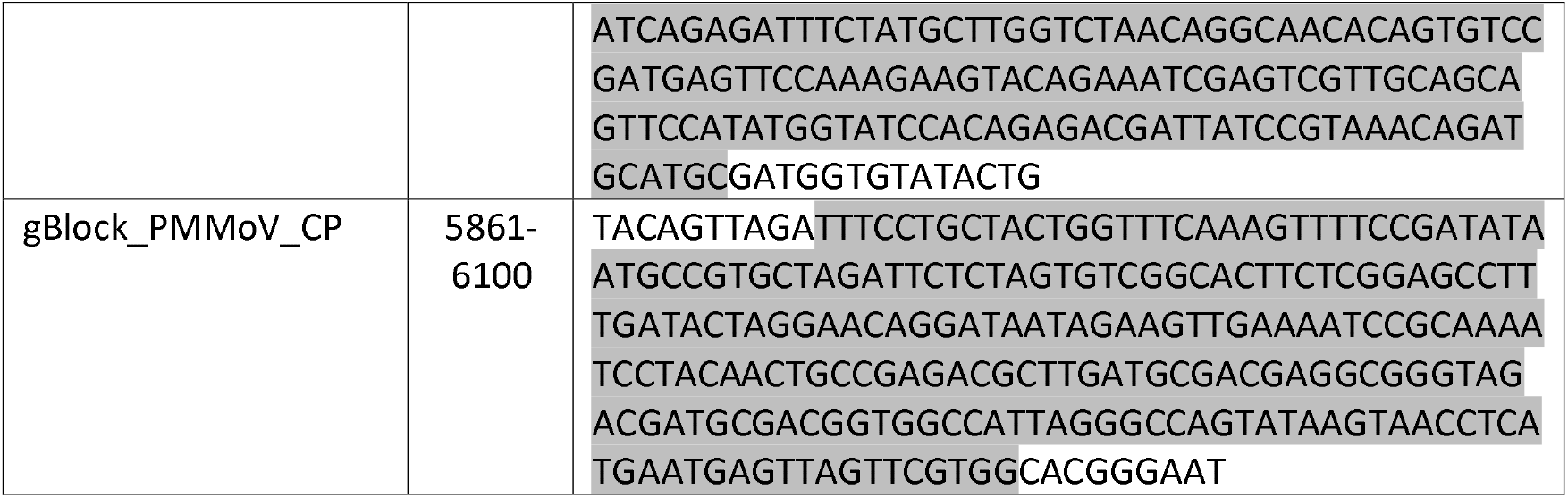
Sequences and positions of the gBlock positive controls for each RT-LAMP assay. Highlighted areas show the section of the genome targeted by each primer set.

### 2.2 RT-LAMP Reaction

Primers for each assay were mixed according to the New England Biolabs (NEB) standard LAMP protocol, utilizing a 10X primer master mix containing 16 μL of the FIP and BIP primers, 2 μL of the F3 and B3 primers, and 4 μL of the LF and LB primers in PCR-grade water (New England Biolabs, 2020). In the PMMoV_CP assay, the 4 μL LF primer was replaced with water in the primer master mix. Two gBlocks™ (IDT, USA) were synthesized to include regions amplified by each primer set (Table 2) and served as positive controls for the analysis.

The RT-LAMP reaction was made using a slightly modified NEB LAMP protocol for a 25 μL reaction volume. In short, the RT-LAMP reaction contains 12.5 μL of WarmStart® Colorimetric LAMP 2X Master Mix with UDG (New England Biolabs, USA), 4.0 μL of PCR-grade water, 2.5 μL of the respective primer mix, and 5.0 μL of a 5% (w/v) solution of pullulan (Sigma-Aldrich, USA) in PCR-grade water. The final 1 μL was reserved for the sample template; in the case of no template control (NTC) samples, 1 μL of PCR-grade water was added to the reaction in lieu of target RNA. The temperature of the reaction was maintained at 62°C for 60 minutes. This assay uses the pH indicator phenol red to detect the pH of the reaction as it changes with the amplification of nucleic acids. A change in color from a pink, negative result to a yellow, positive result signals a successful detection of the target. The reaction progress was monitored by removing the reaction tubes from the incubator at five-minute intervals for the first 30 minutes and then again at the 60-minute endpoint, taking a photograph against a white background, and quickly returning the tubes to the incubator.

During the evaluation of assay performance, two modifications were incorporated into the RT-LAMP reactions to decrease the rate of false positives and inhibit non-specific amplification, both of which are common issues associated with LAMP assays (Alhamid et al., 2023; Gao et al., 2019). The first was the addition of pullulan to the reaction mix. Pullulan is a polysaccharide that has been shown to increase the specificity of the LAMP reaction by inhibiting non-specific amplification (Gao et al., 2019). Second, the reaction temperature was reduced from NEB’s default recommended temperature of 65°C by 3°C to 62°C based on a temperature optimization curve with the pullulan addition and the primer set PMMoV_RdRp (Figure S1).

### 2.3 gBlock Serial Dilutions to Determine Assay Sensitivity

To establish the detection limit of the PMMoV RT-LAMP assay, each gBlock was diluted to a concentration of 10^9^ copies/μL from the stock concentration of 10 ng/μL using the online ThermoFisher DNA Copy Number and Dilution Calculator, and then serially diluted to a concentration of 1 copy/μL. One μL of the gBlock serial dilution was then added to the RT-LAMP reaction and incubated for 60 minutes to evaluate the detection limit for each RT-LAMP assay.

### 2.4 RT-LAMP Specificity Against Related Plant Viruses

To evaluate the specificity of the RT-LAMP reactions, each assay was tested against purified virus RNA from the following related tobamoviruses: tropical soda apple mosaic virus (*Tobamovirus tropici*, TSAMV), TMGMV, TMV, and tomato mosaic virus (*Tobamovirus tomatotessellati*, ToMV). For positive and negative controls, pepper plants (*Capsicum annuum* cv. Aristotle) were grown and maintained in an air-conditioned greenhouse under natural lighting with a daytime high temperature of 30°C. Positive control plants were mechanically inoculated with PMMoV-infected *Nicotiana benthamiana* tissue homogenized in 20 mM sodium phosphate buffer (pH 7.0) containing 1% (wt/vol) Celite. Purified PMMoV RNA from infected pepper plant leaf tissue and total RNA from non-infected pepper plant leaf tissue was extracted using the RNeasy Plant Mini Kit (Qiagen, Germany) and were used as positive and negative controls, respectively. All purified virus RNA and pepper plant total RNA was standardized to 50 ng/μL and 1 μL of RNA was used for each reaction.

## 3. Results

### 3.5 RT-LAMP Assay Detection Limit

The PMMoV_RdRp RT-LAMP reaction began to change color within the first ten minutes of incubation, indicating positive PMMoV detection for concentrations of 10^6^ - 10^9^ copies/μL (Figure 1). The assay reached the detection limit of 100 copies/μL in two of the three replicates and 10 copies/μL in one replicate 20 minutes from the start of incubation. These results were maintained until the end of the 60-minute reaction. The reactions containing one copy of the gBlock and the no-template control (NTC) remained negative throughout the assay.

**Figure 1.**
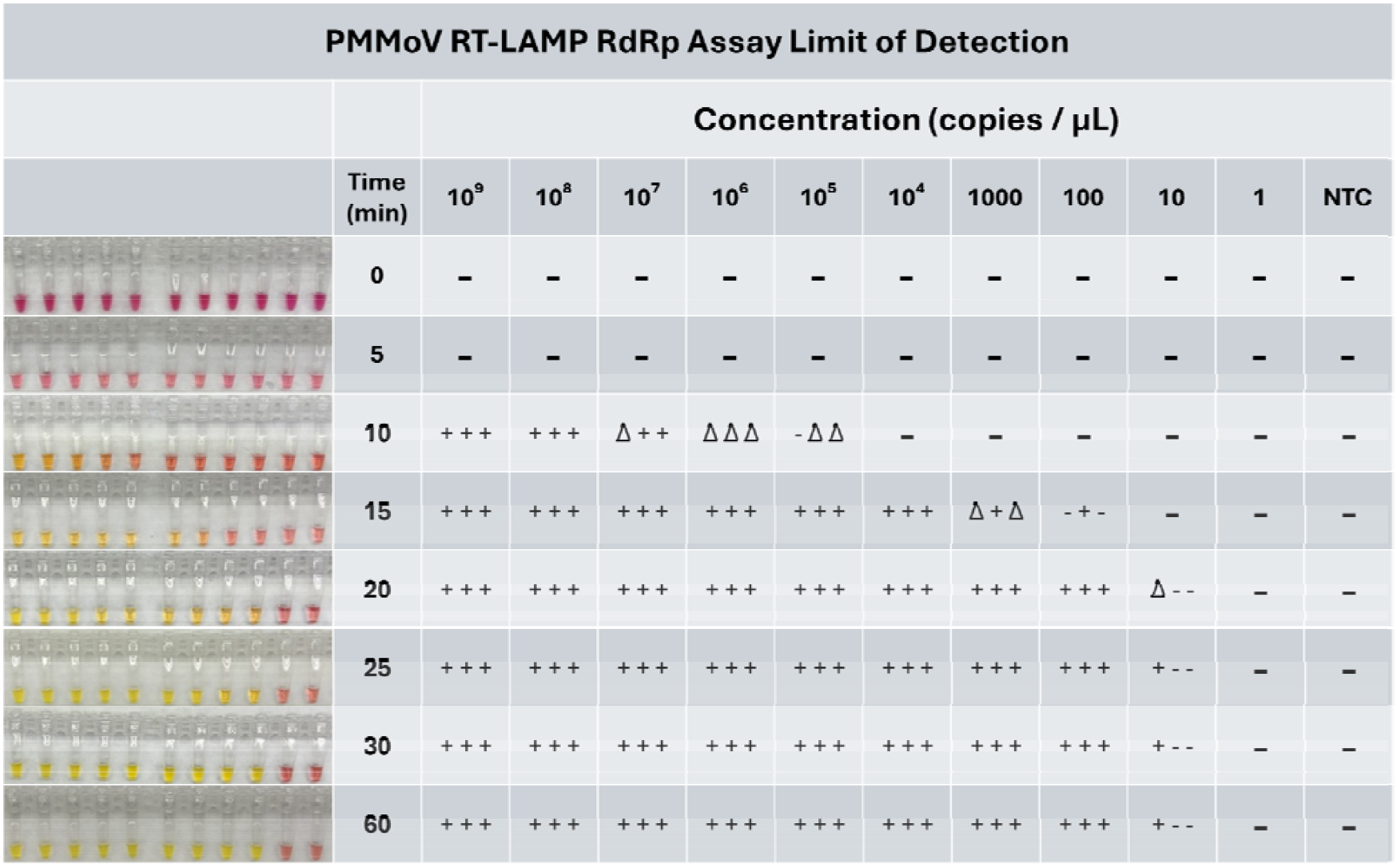
PMMoV_RdRp Assay Limit of Detection Experiment. Photographs of one of the triplicate runs (left) and a table with symbols (“+” = positive result, “Δ” = changing, “—” = negative result) of all triplicate runs (right). The first positive result was observed at t = 10 minutes, and the assay’s detection limit was observed at t = 20 minutes at a concentration of 100 copies/μL. NTC = no template control, using sterile water instead of the gBlock.

The PMMoV_CP RT-LAMP assay detected PMMoV at the highest concentrations tested within twenty minutes (Figure 2). After 25 minutes, the samples containing concentrations between 10^6^ - 10^9^ copies/μL changed colors, indicating a positive result. After 30 minutes, the concentration of 1000 copies/μL turned positive in two replicates and 100 copies/μL in one replicate. There was no further change in color between 30 minutes and 60 minutes. The reactions containing ten and one copy of the gBlock and the NTC remained negative throughout the assay.

**Figure 2.**
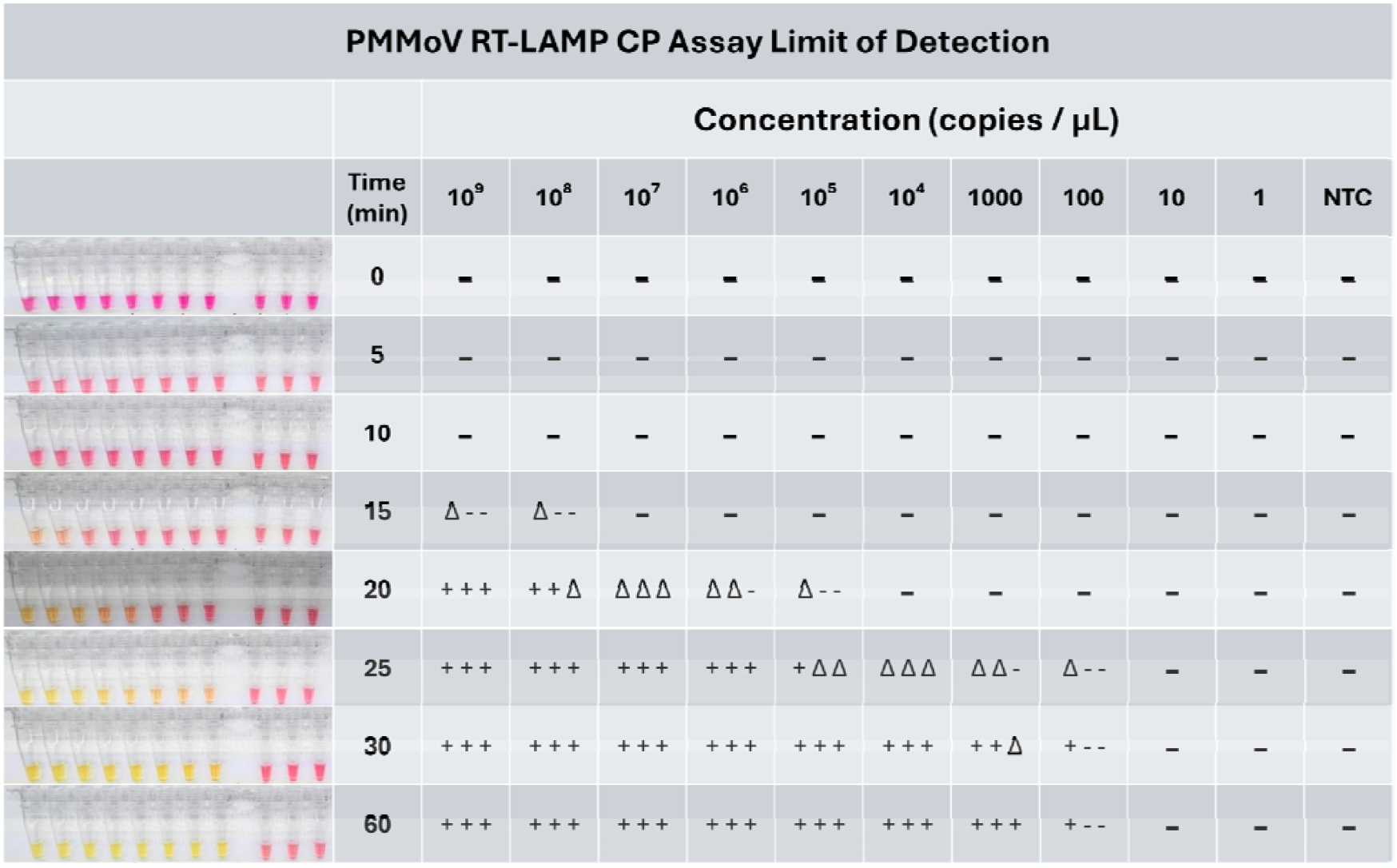
PMMoV_CP Assay Limit of Detection Experiment. Photographs of one of the triplicate runs (left) and a table of all three triplicate results (“+” = positive result, “Δ” = changing, “—” = negative result) at each time point (right). The first positive result was observed at t = 20 minutes, and the detection limit was observed at t = 30 minutes at a concentration of 1000 copies/μL. NTC = no template control, using sterile water instead of the gBlock.

### 3.2 RT-LAMP Assay Specificity

Neither PMMoV RT-LAMP assay demonstrated non-specific amplification of TMV, ToMV, TMGMV, or TSAMV (Figures 3 and 4). Non-infected pepper plant tissue and the NTC also remained negative throughout both experiments. Positive detection of PMMoV RNA occurred at 15 minutes for the PMMoV_RdRp assay (Figure 3) and 20 minutes for the PMMoV_CP assay (Figure 4).

**Figure 3.**
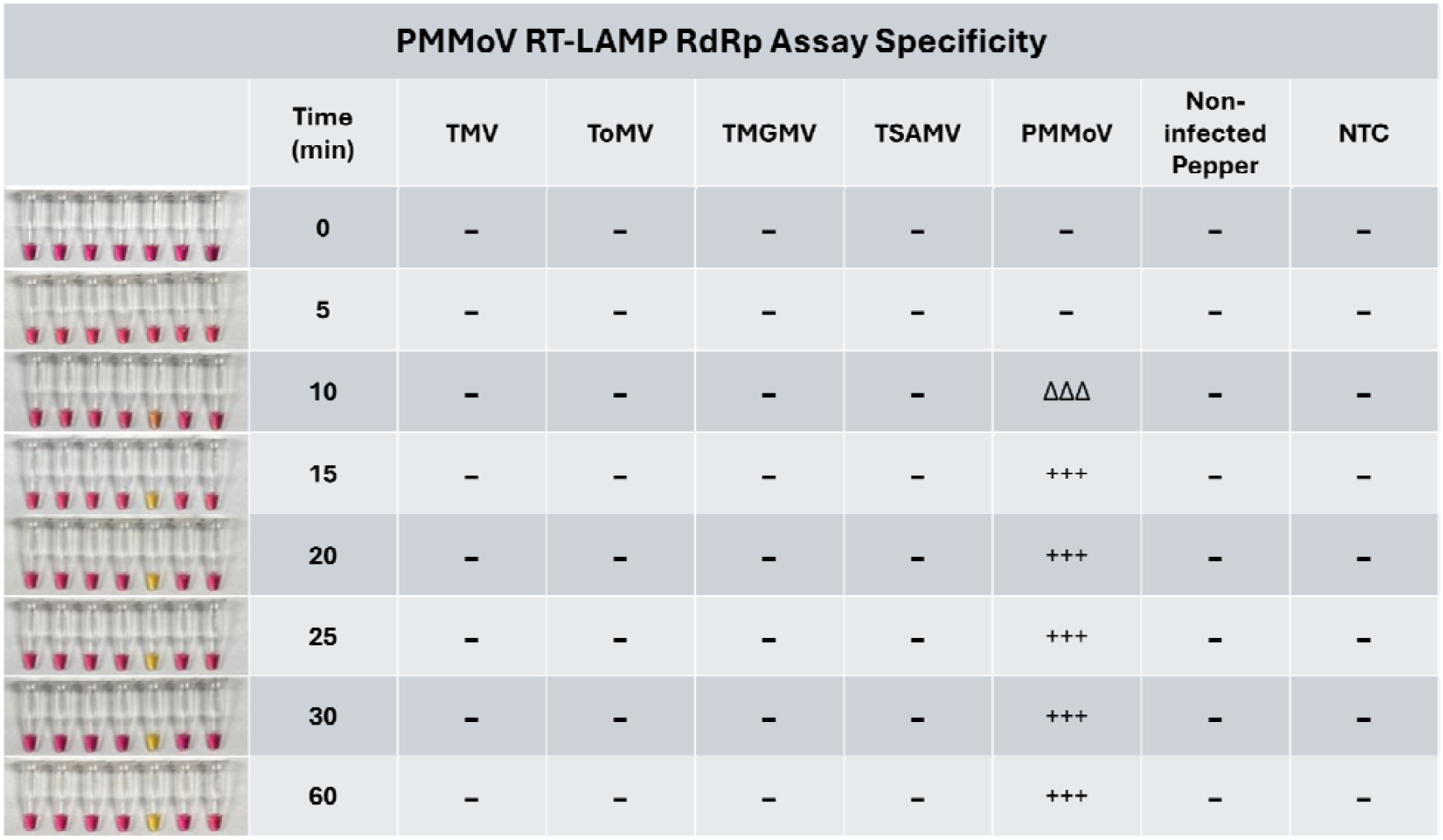
PMMoV_RdRp Assay Specificity Experiment. Photographs of one of the three triplicate runs (left) and a table of all three triplicate results (“+” = positive result, “Δ” = changing, “—” = negative result) at each time point (right). PMMoV RNA was detected at t = 15 minutes in all three triplicates. Related virus RNA from TMV, ToMV, TMGMV, and TSAMV remained negative throughout the assay. Non-infected, RNA-extracted pepper tissue and the NTC (no template control) also remained negative.

**Figure 4.**
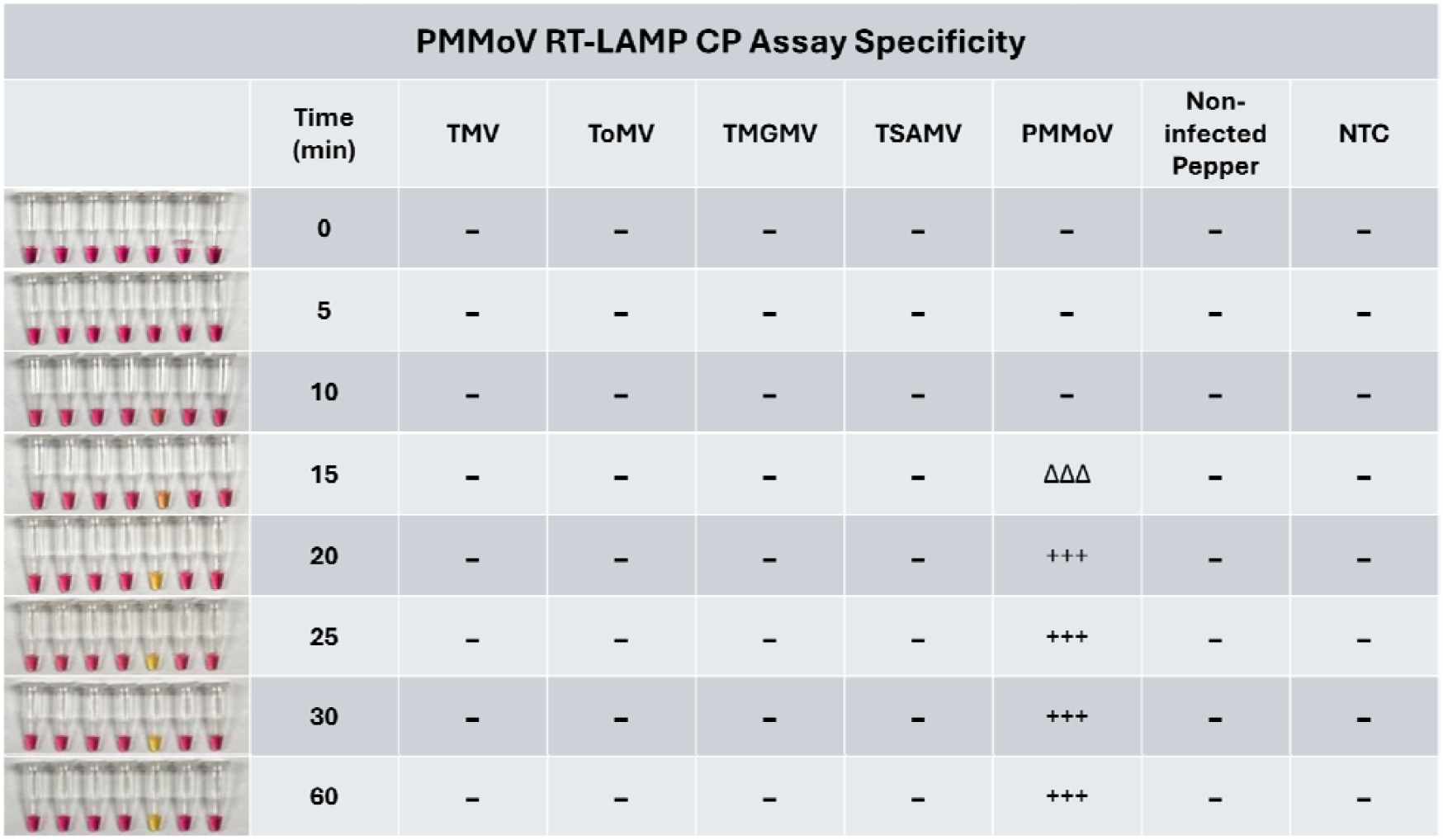
PMMoV_CP Assay Specificity Experiment. Photographs of one of the three triplicate runs (left) and a table of all three triplicate results (“+” = positive result, “Δ” = changing, “—” = negative result) at each time point (right). PMMoV RNA was detected at t = 20 minutes in all three triplicates. Related virus RNA from TMV, ToMV, TMGMV, and TSAMV remained negative throughout the assay. Non-infected, RNA-extracted pepper tissue and the NTC (no template control) also remained negative.

## 4. Discussion

Current methodologies for the detection of PMMoV, such as RT-(q)PCR, ELISA, and immunoassays can have drawbacks, including high costs, requirements for cold storage and specialized laboratory equipment, long times to yield positive results, and varying degrees of specificity and sensitivity. Conversely, the PMMoV RT-LAMP assays described here are sensitive, specific, and inexpensive, providing results at the assay’s detection limit within 20 minutes (RdRp assay) or 30 minutes (CP assay) without needing dedicated laboratory equipment, which makes them suitable for rapid, point-of-use detection. The detection limit of the PMMoV_RdRp assay is comparable to the established RT-qPCR (Rosario et al., 2009; Symonds et al., 2016), detecting as few as 100 copies/μL of the PMMoV gBlock. The detection limit of the PMMoV_CP assay is an order of magnitude higher, at 1,000 copies/μL of the PMMoV gBlock, and therefore less sensitive than the established RT-qPCR assay. Both PMMoV RT-LAMP assays described here are highly specific to PMMoV and do not amplify four closely related tobamoviruses or total RNA from non-infected pepper plant tissue. A recent study described a single nucleotide polymorphism-specific duplex PfAgo RT-LAMP assay for PMMoV and SARS-CoV-2 with a PMMoV detection limit of 21 gene copies per μL on standards (Oh et al., 2024) but did not compare against related tobamoviruses. Future studies should directly compare the sensitivity and specificity of these assays.

Early detection of PMMoV in agricultural settings is crucial to preventing the spread of the virus and the resulting crop loss. Currently, physical observation of greenhouses and crop fields is employed to look for plants showing symptoms of infection (e.g., mottling, mosaic, stunting). Plants showing disease signs must still be tested to identify the causative agent. The RT-LAMP assays described here provide an additional methodology for pathogen confirmation that can be used as an alternative for RT-PCR or ELISA. An expanded toolbox for rapid, point-of-use confirmation of PMMoV in crops will enable early identification of PMMoV infection (Kalimuthu et al., 2022) and the implementation of strategies to prevent further spread (Tatineni and Hein, 2023).

Due to its abundance in wastewater, PMMoV is also used for monitoring wastewater contamination in recreational waters (González-Fernández et al., 2023, 2021; Symonds et al., 2019), shellfish growing waters (Gyawali et al., 2019; Symonds et al., 2017), and waters used for agricultural irrigation (Anderson-Coughlin et al., 2021; González-Fernández et al., 2023; Symonds et al., 2014). PMMoV is used as a process indicator worldwide for wastewater monitoring of human viruses such as norovirus GI and GII, RSV, influenza A, and adenovirus (Eifan et al., 2023; Hughes et al., 2022; Mercier et al., 2022; Miura et al., 2024; Ochar et al., 2023; Parkins et al., 2024). Since the COVID-19 pandemic, PMMoV has also been used to normalize SARS-CoV-2 data to track infections in the populace (Malla et al., 2024; Mercier et al., 2022; Parkins et al., 2024; Wolfe et al., 2021). Additionally, PMMoV has been used as a marker for monitoring wastewater treatment plant efficacy (Iwamoto et al., 2022) and for evaluating emerging wastewater treatment technologies (Symonds et al., 2015). PMMoV detection in these capacities provides valuable insights into the health of human communities, aquatic environments, and the efficiency of wastewater treatment and technologies.

The RT-LAMP assays described here are advantageous for instances where rapid detection of PMMoV is needed or when researchers lack the costly instrumentation and reagents needed for PCR-based assays. Compared to the established PCR and ELISA methods, RT-LAMP provides an equally specific and sensitive assay while maintaining relatively lower costs and yielding rapid results. In addition, the simplicity of an isothermal method means that RT-LAMP could make an effective tool for researchers seeking to detect PMMoV in the field and potentially can be adapted for point-of-use testing. Given the relevance of PMMoV across multiple fields of study, such as agriculture, wastewater treatment, recreational water quality testing, and food safety, many potential uses exist for the RT-LAMP assays described here.

## Declaration of Competing Interests

The authors declare that they have no known competing financial interests or personal relationships that could have appeared to influence the work reported in this paper.

## Acknowledgments

Funding for this work was made available through award NA23NOS4690261 of the NOAA National Centers for Coastal Ocean Science Competitive Research Program to the University of South Florida Tampa Bay Surveillance Project. The authors thank Ian Hewson for his advice on RT-LAMP.

## Supplementary Figure 1

**Figure S1.**
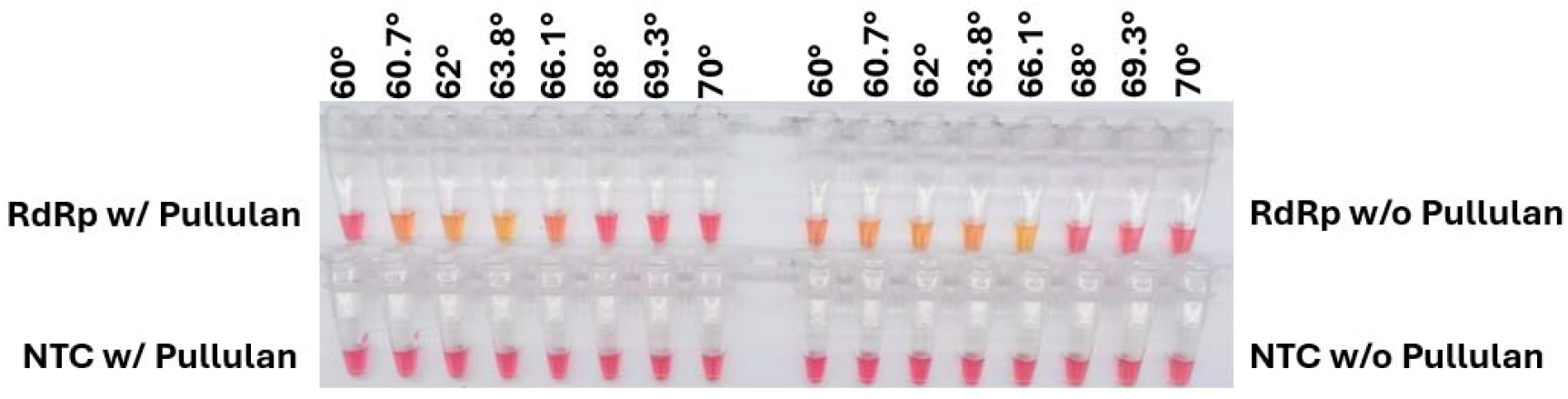
Temperature optimization of the RT-LAMP assay comparing the addition of pullulan vs no addition of pullulan. The experiment used the PMMoV_RdRp assay targeting the PMMoV_gBlock_RdRp at 1000 copies/μL concentration. The photograph above was taken 15 minutes into the reaction at the first sign of a positive result and shows that the optimal temperature of the reaction with the addition of pullulan decreased compared to the standard reaction temperature.

## Notes

### Competing Interest Statement

The authors have declared no competing interest.

### Summary of Updates

Minor revisions have been made to clarify details and a reference to a previously published RT-LAMP assay for PMMoV that was accidentally omitted in the prior version (Oh et al., 2024) has been added.

